# Multiway Canonical Correlation Analysis of Brain Signals

**DOI:** 10.1101/344960

**Authors:** Alain de Cheveigné, Giovanni M. Di Liberto, Dorothée Arzounian, Daniel D.E. Wong, Jens Hjortkjær, Søren Fuglsang, Lucas C. Parra

## Abstract

Brain signals recorded with electroencephalography (EEG), magnetoencephalography (MEG) and related techniques often have poor signal-to-noise ratio due to the presence of multiple competing sources and artifacts. A common remedy is to average over repeats of the same stimulus, but this is not applicable for temporally extended stimuli that are presented only once (speech, music, movies, natural sound). An alternative is to average responses over multiple subjects that were presented with the same identical stimuli, but differences in geometry of brain sources and sensors reduce the effectiveness of this solution. Multiway canonical correlation analysis (MCCA) brings a solution to this problem by allowing data from multiple subjects to be fused in such a way as to extract components common to all. This paper reviews the method, offers application examples that illustrate its effectiveness, and outlines the caveats and risks entailed by the method.

## 1 Introduction

Stimulus-driven signals recorded with electroencephalography (EEG), magnetencephalography (MEG) and related techniques compete with much stronger sources within the brain, the body, and the environment. The signal of interest usually represents only a fraction of the signal power at the electrode or sensor. To overcome the noise and artifacts, a common practice is to present the same stimulus multiple times and average the responses over repeated presentations. Supposing that the response is the same for all presentations, and the noise is un-correlated between presentations, the signal-to-noise power ratio (SNR) improves with the number of repeats. SNR can be further improved by combining signals across sensors, i.e. spatial filtering. Spatial filters can be optimized based on assumptions about signal and noise (de Cheveigné and Parra, 2014), and this combination of temporal averaging and spatial filtering can greatly improve the SNR. However, averaging and optimization are not applicable if the stimulus is presented only once, for example because it is too long to be repeated (e.g. a long sample of speech or music), or because one wishes to probe a phenomenon likely to fade with repetitions (e.g. surprise).

Instead of presenting the same stimulus multiple times to one subject, one can also present the same stimulus to multiple subjects just once. To the extent that different subjects’ brains are functionally similar, we expect similar responses (Hasson et al., 2004; Dmochowski et al., 2012; Lankinen et al., 2014). Unfortunately, the position or orientation of neural sources relative to sensors or electrodes is likely to differ across subjects, so averaging over subjects in sensor space is suboptimal. In order to compare between subjects, or average over subjects, we first need some way to transform the data of each to a common representation that is comparable across subjects. This can be accomplished with spatial filters that are tuned to each individual subject (e.g. Haxby et al., 2011; Lankinen et al., 2014).

Canonical Correlation Analysis (CCA) is a powerful technique to find linear components that are correlated between two data matrices (Hotelling, 1936). Given two matrices **X**_1_ and **X**_2_ of size *T* × *d*_1_ and *T* × *d*_2_, CCA produces transform matrices **V**_1_ and **V**_2_ of sizes *d*_1_ × *d*_0_ and *d*_2_ × *d*_0_, where *d*_0_ is at most equal to the smaller of *d*_1_ and *d*_2_. The columns of **Y**_1_ = **X**_1_ **V**_1_ are of norm 1 and mutually uncorrelated between each other, as are the columns of **Y**_2_ = **X**_2_**V**_2_, while, more importantly, corresponding columns from each (“canonical correlate pairs”) are maximally correlated. The first pair of canonical correlates (CC) defines the linear combinations of each data matrix with the *highest possible correlation* between them. The next pair of CCs defines the most highly correlated combination that is uncorrelated from the first pair, and so-on. Applied to data from two subjects, CCA can find spatial filters that maximize the brain activity common to both, transforming both subject’s data so that they can more easily be compared or averaged. However, CCA does not address the issue of comparing or merging responses across more than two subjects.

Extensions to connect multiple data matrices have been proposed under names such as *multiple CCA* (Gross and Tibshirani, 2015; Witten and Tibshirani, 2009), *multiway CCA* (Sturm, 2016; Zhang et al., 2011), *multiset CCA* (Takane et al., 2008; Correa et al., 2010b,a; Hwang et al., 2012; Lankinen et al., 2014; Zhang et al., 2017; Via, Javier, Ignacio Santamaria and Pérez, 2005; Li et al., 2009), or *generalized CCA* (Kiers et al., 1994; Afshin-Pour et al., 2012; Melzer et al., 2001; Tenenhaus, 2011; Tenenhaus et al., 2015; Velden, 2011; Fu et al., 2017). This diversity in names covers a diversity of formulations (Kettenring, 1971) that all share the aim of finding components that are similar across data matrices. Recent progress addresses regularization (Tenenhaus, 2011), sparsity (Fu et al., 2017; Tenenhaus et al., 2015), missing data (van de Velden and Takane, 2012), nonlinearity (Melzer et al., 2001), or deep learning (Benton et al., 2017). Using similar techniques, independent Component Analysis (ICA) has been generalized under the name of group ICA (GICA) (Eichele et al., 2011; Calhoun and Adali, 2012; Huster et al., 2015; Huster and Raud, 2018).

CCA has been used extensively for brain data analysis and modality fusion (Sui et al., 2012; Dähne et al., 2015; Dmochowski et al., 2017), and several studies have applied multiway CCA (MCCA) and variants thereof to merge data across subjects (Correa et al., 2010b; Afshin-Pour et al., 2012, 2014; Lankinen et al., 2014; Zhang et al., 2017; Li et al., 2009; Hwang et al., 2012; Karhunen et al., 2013; Haxby et al., 2011; Lankinen et al., 2014; Sturm, 2016; Zhang et al., 2017; Lankinen et al., 2018). This paper builds on those studies with the aim to better understand the range of applicability of the tool, what is achieved, and what are the caveats. We describe a simple formulation of MCCA that is easy to understand and explain.

We show that MCCA can be applied effectively to multi-subject datasets of EEG or fMRI, both to *denoise* the data prior to further analyses, and to *summarize* the data and reveal traits common across the population of subjects. MCCA-based denoising yields significantly better scores in an auditory stimulus-response classification task, and MCCA-based joint analysis of fMRI data reveals detailed subject-specific activation topographies. The aims of this paper are (a) to provide an intuitive understanding of MCCA, (b) investigate ways in which it can be put to use, and (c) demonstrate its effectiveness for a range of common tasks in the analysis of brain data.

## 2 Methods

In this section we describe a simple formulation of MCCA, show how it can be applied to a variety of tasks, and give details of the real and synthetic data sets used by the examples reported in the Results.

### 2.1 Data analysis

#### Signal model

Assume a data set consisting of *N* data matrices, each comprised of a time series matrix **X***_n_* of dimensions *T* (time) × *d_n_* (channels). These could represent EEG, MEG or fMRI data recorded from N different subjects in response to the same stimulus. They could also be data from multiple imaging modalities gathered from the same subject. Each matrix **X***_n_* consists of linear combinations of a set of sources S common to all data matrices, to which is added a “noise” matrix **N***_n_* of sources uncorrelated with S, and uncorrelated with the noise matrices **N**_*n*′≠*n*_ added to the other data matrices:

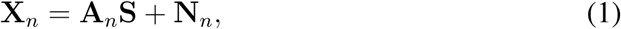

where A*_n_* is a mixing matrix specific to subject *n*. The sources S might represent brain sources or networks driven by the same stimulus similarly across different subjects. We are interested in finding these “shared sources” and suppressing the noise. Note that this model assumes that responses of different subjects share the same source *time course*, but not necessarily the same spatial pattern over channels. The assumption of uncorrelated noise is usually only approximately met, due to spurious correlations.

#### A simple CCA formulation

Consider two data matrices, **X**_1_ and **X**_2_ of size *T* × *d* where *T* is time and *d* the number of channels. All data are assumed to have zero mean. Each matrix is spatially whitened by applying principal component analysis (PCA) and scaling each principal component (PC) to unit norm to obtain whitened matrices 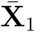 and 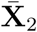. Whitened data are then concatenated and submitted to a new PCA to obtain a matrix **Y** = [**X**_1_, **X**_2_] V of size *T* × 2*d*, where **V** combines the whitening and second PCA matrices (Fig. 1 left). The submatrices **V**_1_ and **V**_2_ formed of the first and last *d* rows of **V** define transforms applicable to each data matrix:

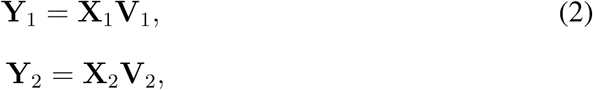

with **Y** = **Y**_1_ + **Y**_2_ (Fig. 1 center).

**Figure 1:**
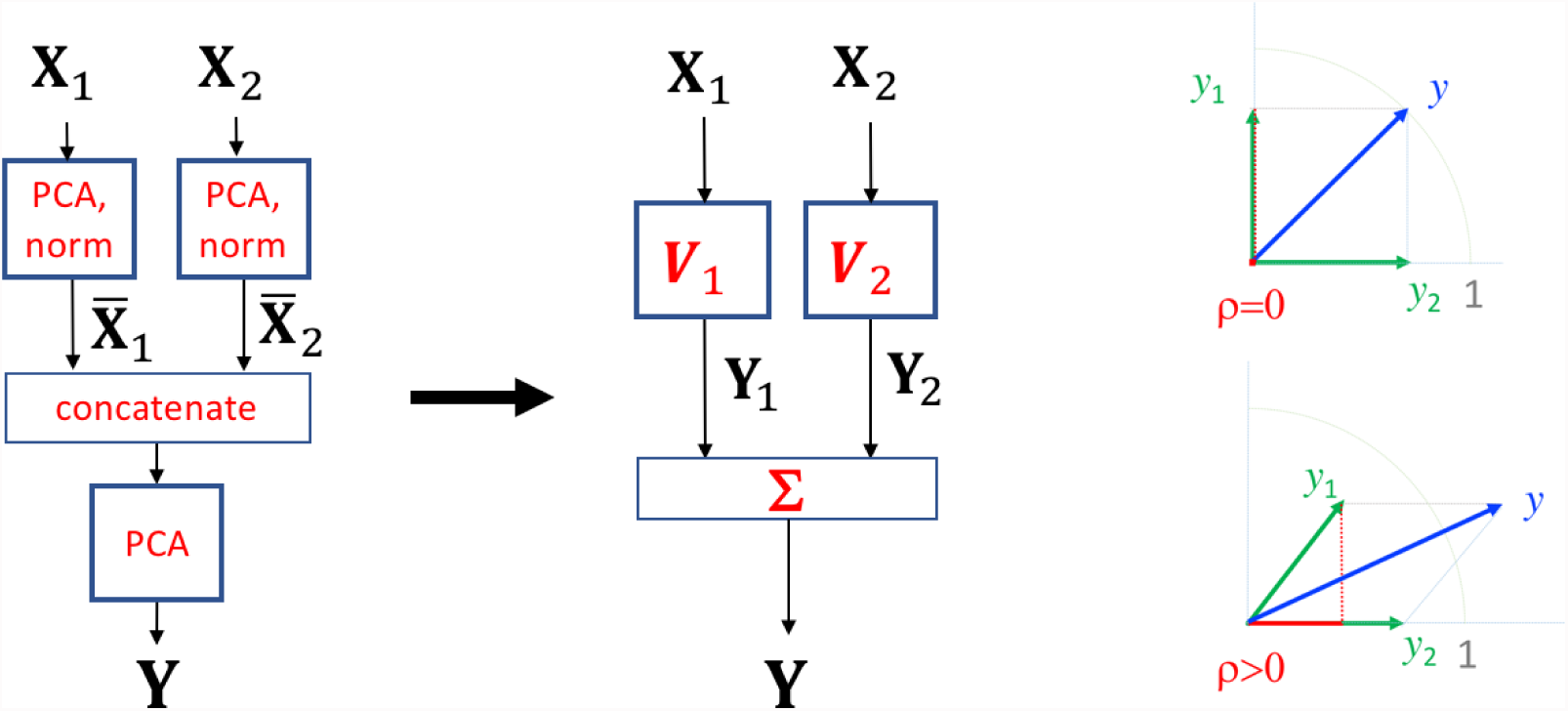
Block diagram of the simple CCA formulation. Left: each data matrix is whitened by PCA followed by normalization. Normalized PCs from both data matrices are concatenated side by side and submitted to a final PCA. Center: the matrix ***Y*** of summary components (SC) can be expressed as the sum of individual transforms ***Y***_1_ = ***X***_1_***V***_1_ and ***Y***_2_ = ***X***_2_***V***_2_ (canonical correlates, CC). The transforms ***V***_1_ and ***V***_2_ combine the whitening and PCA matrices. Right: rotating vectors y_1_ and y_2_ to maximize the norm of their sum is equivalent to maximizing their correlation coefficient ρ symbolized by the projection of y1 on y2 (red line).

The outcome of this analysis is equivalent to standard CCA, as explained in the Discussion, the first *d* columns of **Y**_1_ and **Y**_2_ forming canonical pairs (within a scaling factor). Indeed, rotating 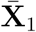 and 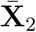 to maximize the correlation of the resulting **Y**_1_ and **Y**_2_, as required by the CCA objective, is equivalent to rotating with the goal of maximizing the norm of their sum, **Y**_1_ + **Y**_2_, as achieved by the second PCA (Fig. 1 right). The appeal of this formulation is that it is easily extendable to multiple data matrices.

#### A simple MCCA formulation

Consider *N* data matrices **X***_n_* each of size *T* × *d* with zero mean. Each data matrix is spatially whitened by applying PCA and scaling all PCs to unit norm to obtain whitened matrices 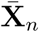. Whitened data are then concatenated along the component dimension and submitted to a second PCA to obtain a matrix **Y** = [**X**_1_ … **X***_N_*] **V** of size *T* × *D*, *D* = *Nd*, where **V** combines the whitening and second PCA matrices (Fig. 2 left). The submatrices **V***_n_* of **V** of size *d* × *D* formed by extracting successive d-row blocks of **V** define transforms applicable to each data matrix:

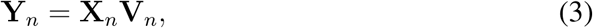

with 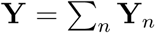 (Fig. 2, right). If data matrices have different numbers of channels *d_n_*, then **V***_n_* has size *d_n_* × *D* where 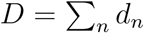. We call the columns of **Y***_n_ canonical correlates* (CCs) by analogy with CCA, and those of **Y** *summary components* (SC). Each SC is a sum of CCs over data sets. Columns of **Y** are mutually orthogonal by virtue of the final PCA, but the same is not usually true of **Y***_n_*. With *D* > *d* columns, **Y***_n_* forms an *overcomplete basis* of the patterns spanned by **X***_n_*. This formulation of MCCA is equivalent to the SUMCORR formulation of Kettenring (1971) as explained in the Discussion (Parra, 2018). The appeal of this formulation is that it is conceptually and computationally straightforward. PCs can be discarded from the initial PCAs, so as to control dimensionality and limit overfitting effects (next section).

**Figure 2:**
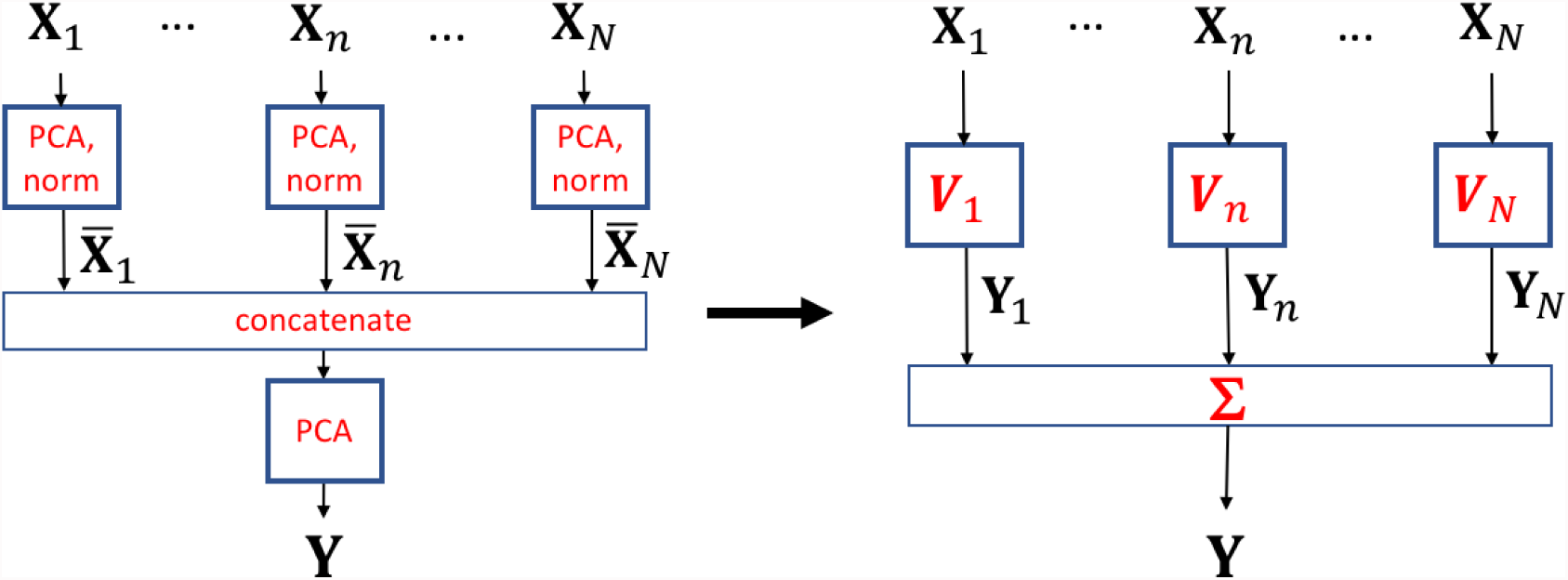
Block diagram of the simple MCCA formulation. Left: each data matrix ***X***_n_ is whitened by PCA followed by normalization. Normalized PCs from all data matrices are concatenated side by side and submitted to a final PCA. Right: the matrix ***Y*** of summary components (SC) can be expressed as the sum of individual transforms ***Y***_n_ = ***X***_n_***V***_n_ (canonical correlates, CC).

The variances of the summary components (the columns of **Y**) reflect the degree to which temporal patterns are shared between data matrices (Fig. 3) – the variance of each SC corresponding to the degree of correlation of each shared dimension found in the data. If the data matrices **X***_n_* share no components, the variances of all SCs are one (Fig. 3 a). If a component is shared by all *N* data matrices, the norm of the first SC is *N* (Fig. 3 d). For data matrices with a small number of samples, spurious correlations may cause the variance profile to be skewed (Fig. 3 b). In real data, shared activity often shows up as components with variance elevated relative to this background (Fig. 3 c).

**Figure 3:**
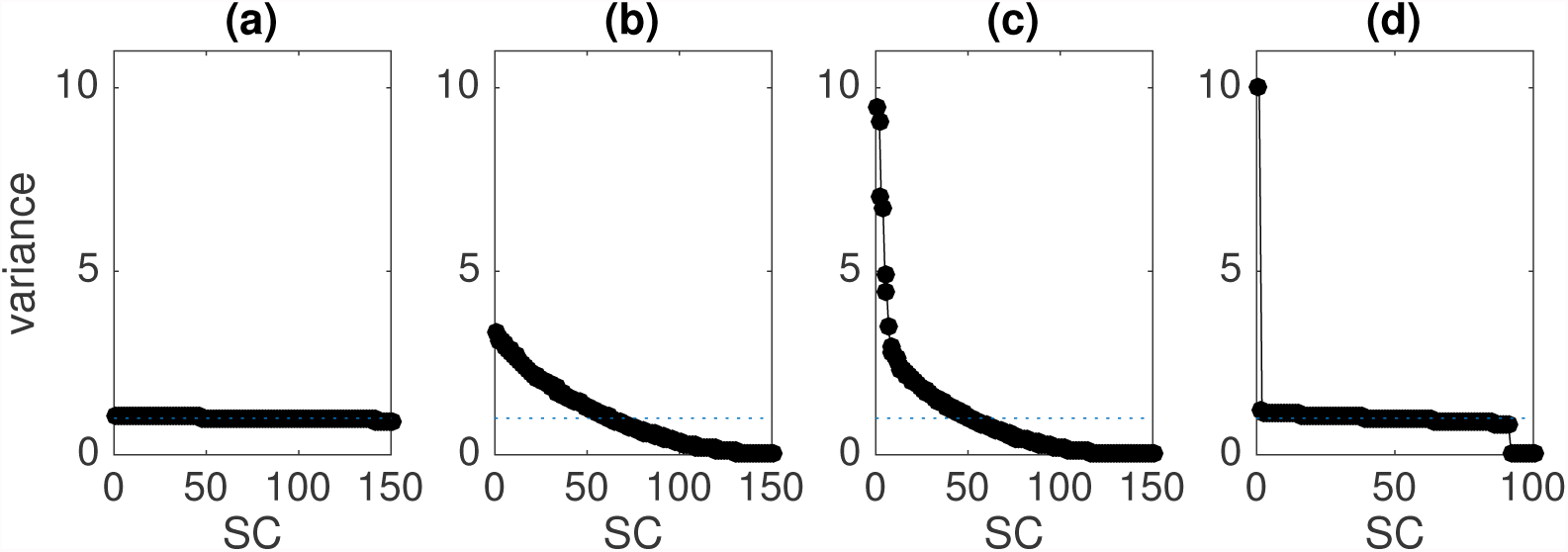
Behavior of the SC variance as a function of order for MCCA analyses applied to 4 different types of dataset, each involving 10 data matrices. (a) Each data matrix consisted of an independent 10000 × 15 matrix of Gaussian white noise. In this case the SC variance profile is flat since there is no (or little) correlation between data matrices. (b) Each data matrix consisted of a 165 × 15 matrix of independent and uncorrelated Gaussian noise. In this case the SC variance profile is skewed, reflecting spurious numerical correlations between the statistically independent columns. (c) Each data matrix consisted of a 165 × 15 matrix of values derived from fMRI responses of 10 subjects in response to 165 sounds. Prior to MCCA the 6309 voxels were reduced to 15 channels using SVD (see description of Example 6 in the Methods). (d) Each data matrix consisted of a 10000 × 10 matrix of Gaussian white noise with an embedded sinusoid (Example 1, Fig. 4) that was the same in all data matrices. In the last two examples, only a small subset of the MCCA components reflect shared activity as evident by the low SC variance at higher MCCA orders.

#### Reduced-rank MCCA

It is often convenient to reduce the rank of each data matrix 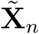 to 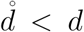 by discarding PCs with smallest variance after the initial PCA. The MCCA transform matrices **V***_n_* are then of size 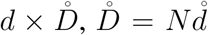, and the CC and SC matrices of size 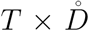. This serves as a form of reglarization that avoids computational issues with rank-deficient data, reduces the risk of overfitting, and limits computation and memory requirements. Importantly, this approach preserves the constraint that the resulting SCs are uncorrelated (Parra et al., 2018).

#### Dealing with data matrices with more channels than samples

CCA fails if the data matrices have fewer samples than channels (*T* ≤ *d*), as is typically the case for fMRI or calcium imaging data for which there are many more voxels or pixels than observation samples (Asendorf, 2015). A simple solution is to replace each data matrix **X***_n_* (size *T* × *d*) by a matrix 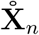 of size 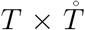 with 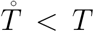 columns that capture the principal temporal patterns spanned by **X***_n_*. This can be done by applying singular value decomposition (SVD) to express the data as

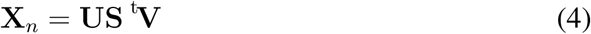

and setting 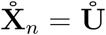 where 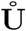 consists of the first 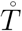 columns of U. Since the 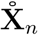 have more samples than channels there is no obstacle to applying MCCA to them. This sequence of operations can be represented by a set of transform matrices **V***_n_* of size 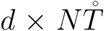. Applying them to the data yields canonical correlate and summary matrices of size 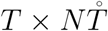. Using this approach, it is straightforward to apply MCCA to datasets with a large number of “channels” such as data from calcium imaging or fMRI. An alternative to SVD is to apply PCA to ^t^**X**_*n*_ and use a subset of the matrix of projection vectors to form 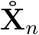, a useful option if **X***_n_* is too large to fit in memory (the required covariance matrix can be calculated in chunks).

### 2.2 Applications of MCCA

#### Quantifying correlation between *N* data matrices

The variance of each column of **Y** indicates the degree to which a component is shared across data matrices. The value is 1 if the data matrices are perfectly uncorrelated, and *N* if all data matrices include that component (Fig. 3). The profile of variances over SCs thus offers a measure of “sharedness” between data matrices (but see Caveats).

#### Summarizing a set of data matrices

The first few columns of 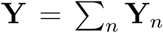 represent temporal patterns that capture most of the correlation across data matrices **X***_n_*. They form a basis of the signal subspace that contains those shared patterns.

#### Denoising

Each data matrix **X***_n_* may be denoised by projecting it to the over-complete basis of CCs, selecting the first 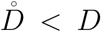 components, and projecting back. We refer to this procedure as “denoising”, as it can be used to attenuate components that are least shared across subjects. This can be summarized by a denoising matrix **D***_n_* product of the first 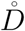 columns of **V***_n_* by the first 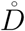 rows of its pseudoinverse. The denoised data are obtained as 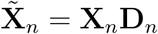.

#### Dimensionality reduction

Dimensionality reduction is often performed by applying PCA to a data matrix and truncating the PC series (Cunningham and Yu, 2014). However, this equates relevance to variance, which may not be appropriate because noise sources can have high variance and useful targets small variance. MCCA can be used to weight dimensions according to their *consistency across data matrices*, which may be a better criterion than variance.

#### Outlier detection

Temporally-local glitches and artifacts may interfere with data interpretation and analysis. Analysis algorithms based on least-squares are particularly sensitive to high-amplitude artifacts. MCCA can be used to derive a cross-subject ‘consensus’ response, so that individual subject’s data points that deviate greatly from the consensus can be flagged as outliers and excluded from analysis.

### 2.3 Details of the evaluation examples

The methods are evaluated using six datasets, including synthetic data, EEG, and fMRI.

#### Example 1 - sinusoidal target in separable noise

Synthetic data for this example consisted of 10 data matrices, each of dimensions 10000 samples × 10 channels. Each was obtained by multiplying 9 Gaussian noise signals (independent and uncorrelated) by a 9 × 10 mixing matrix with random coefficients. To this background of noise was added a “target” consisting of a sinusoidal time series (Fig. 4, left) multiplied by a 1 × 10 mixing matrix with random coefficients. The target was the same for all data matrices, but the mixing matrices differed, as did the noise sources. The SNR was set to 10^−20^, i.e. a very unfavorable SNR, but because the noise is not of full rank the target and background are in principle linearly separable.

**Figure 4:**
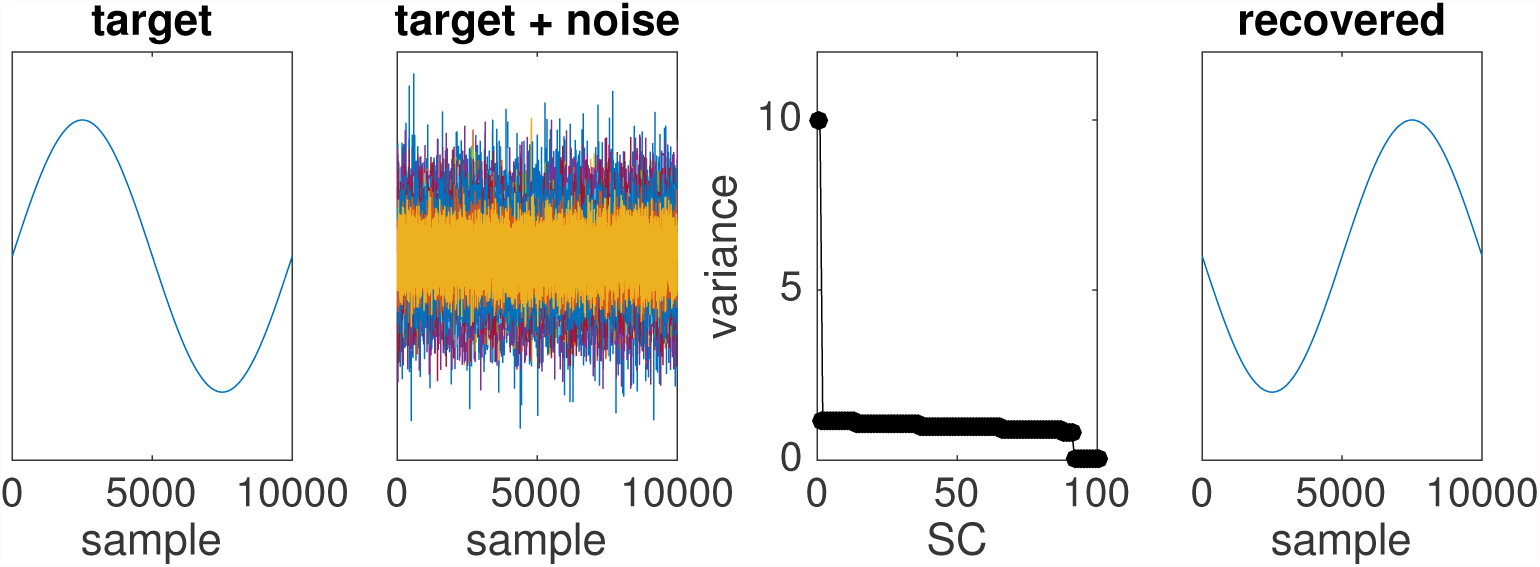
Simulation with separable noise. Left: target signal. Next to left: target in noise at SNR=10^−20^. Next to right: variance of SCs as a function of order. The variance of the first SC is equal to 10 as target is perfectly shared across subjects and mixed in separable noise. Right: target recovered by MCCA (with arbitrary sign).

#### Example 2 - sinusoidal target in non-separable noise

Synthetic data for this example consisted of 10 matrices of dimensions 10000 samples × 10 channels, each obtained by multiplying 10 Gaussian noise sources (independent and uncorrelated) by a 10 × 10 mixing matrix with random coefficients. To this background was added a sinusoidal target as in the previous example, with SNR varied as a parameter. The noise here is full rank so the target and background are not linearly separable.

#### Example 3 - sinusoidal target in EEG noise

Data for this example used EEG to simulate realistic neural activity as background noise. EEG data were recorded during approximately 20 minutes from one subject in the absence of any task, from 40 electrodes (32 standard positions plus additional electrodes on forehead and temple) at 2048 Hz sampling rate with a BioSemi system. A robust polynomial detrending routine (de Cheveigné and Arzounian, 2018) was used to remove slow drifts. Ten “data matrices” were produced by selecting three-second intervals of EEG data with random offsets, removing their means, and adding a target consisting of 4 cycles of a 4 Hz sinusoid multiplied by a 1 × 40 mixing matrix with random coefficients, renewed for each data matrix. The SNR of the target was varied as a parameter.

#### Example 4 - EEG response to tones

Data for this example were borrowed from a study on auditory attention (Southwell et al., 2017). EEG data were recorded using a 64-channel EEG system in response to 120 repetitions of a 1 kHz tone pip with interstimulus interval (ISI) randomized between 750 and 1550 ms (recorded for the purpose of locating electrodes responsive to sound). Data from a subset of 10 subjects were detrended using a robust detrending routine, bad channels were interpolated using spherical interpolation (EEGLAB), and the data were filtered between 2-45 Hz. A peristimulus epoch of duration 1.2 s (starting 0.2 s prestimulus) was defined for each trial, and the corresponding data were extracted as a 3D matrix of dimensions time × channel × trial. For each channel, the 0.2 s prestimulus waveform was averaged over trials and subtracted from that channel’s waveform (“baseline correction”). After applying the first PCA (of the two-step MCCA) to each subject, the first 30 PCs were retained and the remainder discarded.

Two analyses were performed on these data to try to extract the cortical response to the 1 kHz tone from the background EEG noise. In the first, repetition over trials was exploited to design a spatial filter for each subject using the joint diagonalization algorithm (JD) that maximizes the ratio of trial-averaged variance to total variance (de Cheveigné and Simon, 2008; de Cheveigné and Parra, 2014). This resulted in a set of 10 analysis matrices of size 64 × 30, one for each subject. In the second analysis, MCCA was applied, using 30 PCs from each subject in the first PCA, resulting in 10 subject-specific analysis matrices of size 64 × 300.

For each subject, the first column of the JD analysis matrix defines the best linear combination of channels to maximize repeat-reliability across trials, while the first column of the MCCA analysis matrix defines the best linear combination of channels to maximize correlation with the other subjects.

#### Example 5 - EEG response to speech

Data for this example were taken from a study on auditory cortical responses to natural speech (Di Liberto et al., 2015). The same data were also used in a recent study on the application of CCA to speech/EEG decoding (de Cheveigné et al., 2018). We borrowed the data from the first study, and the decoding methods and evaluation metrics from the second, with the purpose of evaluating the benefit of introducing a denoising stage based on MCCA before the speech/EEG decoding stage.

In brief, EEG data were recorded from 8 subjects using a 128-channel BioSemi system with standard electrode layout, at 512 Hz sampling rate. Each subject listened to 32 speech excerpts, each of duration 155 s, from an audio book, presented diotically via headphones, for a total of approximately 1.4 hours. The database included both the audio stimuli and the EEG responses. Further details about the stimulus and recording are available in Di Liberto et al. (2015). The EEG were preprocessed (downsampling to 64 Hz, detrending, artifact removal), and the stimulus temporal envelope calculated as described in de Cheveigné et al. (2018).

A decoding model (de Cheveigné et al., 2018; Dmochowski et al., 2017) was evaluated according to several metrics: correlation, d-prime, and percent-correct classification scores for a match vs mismatch classification task. The classification task consisted in deciding whether a segment of EEG matched the segment of stimulus of same duration that produced it (match) or some unrelated segment (mismatch). The duration of the segment was varied as a parameter from 1 to 64 s.

This task is related to that of determining which of two concurrent voices is the focus of a listener’s attention (cocktail party phenomenon) (Ding and Simon, 2012; Fuglsang et al., 2017; Lalor et al., 2009; Khalighinejad et al., 2017; Koskinen and Seppä, 2014; Martin et al., 2014; Mesgarani and Chang, 2012; Mirkovic et al., 2015; O’Sullivan et al., 2014; Tiitinen et al., 2012; Zion Golumbic et al., 2013), of potential use for the “cognitive control” of an external device such as a hearing aid. The decoding model used CCA to relate the stimulus to the EEG response, producing multiple stimulus-response CC pairs that were used for discrimination. Further details of the decoding model, classification task, and metrics can be found in de Cheveigné et al. (2018). Here, we are only interested in knowing if scores for single-source decoding are improved by introducing a stage of EEG denoising based on MCCA.

For this denoising, the EEG data of each subject were submitted to MCCA, keeping 40 PCs in the first PCA, resulting in a 128 × 320 analysis matrix for each subject. The first 110 columns of this matrix were multiplied by the first 110 rows of its pseudoinverse to yield a 128 × 128 subject-specific denoising matrix. This has the effect of attenuating activity that is *least* correlated with the other subjects.

#### Example 6 - fMRI response to natural sounds

Data for this example were taken from a study that measured fMRI responses to natural sounds (NormanHaignere et al., 2015). Responses were gathered from 10 subjects to each of 165 sounds belonging to 11 categories including speech, music, animal vocalizations, and others. For each subject, the recording session was repeated either twice or 3 times. See Norman-Haignere et al. (2015) for further details. For the present analysis, data for each subject were averaged over repeats and organized as a matrix **X***_n_* of 165 sounds × 6309 voxels (voxels from both hemispheres were used, and voxels outside a subject-specific region of interest that included primary and secondary auditory cortex were set to zero). In this analysis we are interested in finding particular profiles of response over sounds (for example speech vs non-speech, or music vs non-music) and also the brain areas associated with such profiles in each subject.

As there are more “channels” (voxels) than samples (*T* < *d*), an SVD was used as described in the Methods and the first 10 dimensions were used for MCCA. The columns of 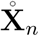 are white so the first PCA can be dispensed of. Matrices 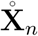 were concatenated and subjected to the second-step PCA of the MCCA algorithm, and the 15 first columns (arbitrary number) of the SC matrix were selected as a basis spanning the profiles over sounds that were most similar across subjects.

To find profiles specific to particular sound categories (e.g. speech, music, etc.), Joint Decorrelation (de Cheveigné and Parra, 2014) was used to find a linear transform applicable to the 15-column basis to maximize the variance over the selected category, relative to the other categories. This can be seen as a rotation of the basis so as to isolate activity specific to processing of that sound category. This 165 × 1 activation profile was then cross-correlated with the 165 × 6309 matrix of fMRI response data of each subject to find the topography specific to that subject (Haufe et al., 2014).

## 3 Results

The MCCA method is evaluated first with synthetic data to get an understanding of its basic properties and capabilities, and then with real EEG and MEG data to see whether these extend to situations of practical use.

### 3.1 Synthetic data

#### Example 1 - sinusoidal target in separable noise

The data consist of 10 matrices made up of a sinusoidal target (Fig. 4, left) common to all data matrices, with added noise distinct across matrices (see Methods). At the unfavorable SNR of 10^−20^ the target is not visible in the raw signal of any of the data matrices (Fig. 4 center), and it cannot be extracted by averaging because of the extremely low SNR and the fact that the mixing coefficients are of random sign. Since the data are separable (the rank of the noise is only 9), the target *can* be recovered by applying the appropriate demixing matrix (inverse of the mixing matrix), however that matrix is unknown.

MCCA applied to the dataset produced projection matrices **V**_*n*_ that recover the target from **X***_n_* (Fig. 4 right). This benefit is similar to that of methods that leverage multiple repetitions to blindly discover spatial filters to improve SNR (de Cheveigné and Simon, 2008; de Cheveigné and Parra, 2014), but instead of repetitions, MCCA leverages the fact that the same target is mixed into multiple data matrices. To summarize, MCCA can reveal a target common across data matrices despite an extremely unfavorable SNR.

#### Example 2 - sinusoidal target in non-separable noise

Data are the same as in the previous example, except that the noise is full rank (10 independent sources mixed in 10 channels) so the target is no longer linearly separable, and one cannot expect to recover the target perfectly, especially at extremely low SNRs. Nonetheless, at a moderately unfavorable SNR (10^−2^ in power) MCCA can recover an estimate of the target that is noisy (Fig. 5 center) but much cleaner than the raw data (not shown). Figure 5 (right) shows the proportion of residual noise in the signal recovered by MCCA as a function of SNR, together with the same proportion for the best raw channel. MCCA provides a clear benefit over a range of SNRs. Two factors can contribute to failure: non-separability per se, and the fact that the algorithm fails to find the ideal demixing matrix. Figure 5 (right) also shows the proportion of residual noise for the ideal demixing matrix (yellow). The MCCA-derived matrix performs only slightly less well than the ideal matrix. To summarize, MCCA is of use even if the data are not separable.

**Figure 5:**
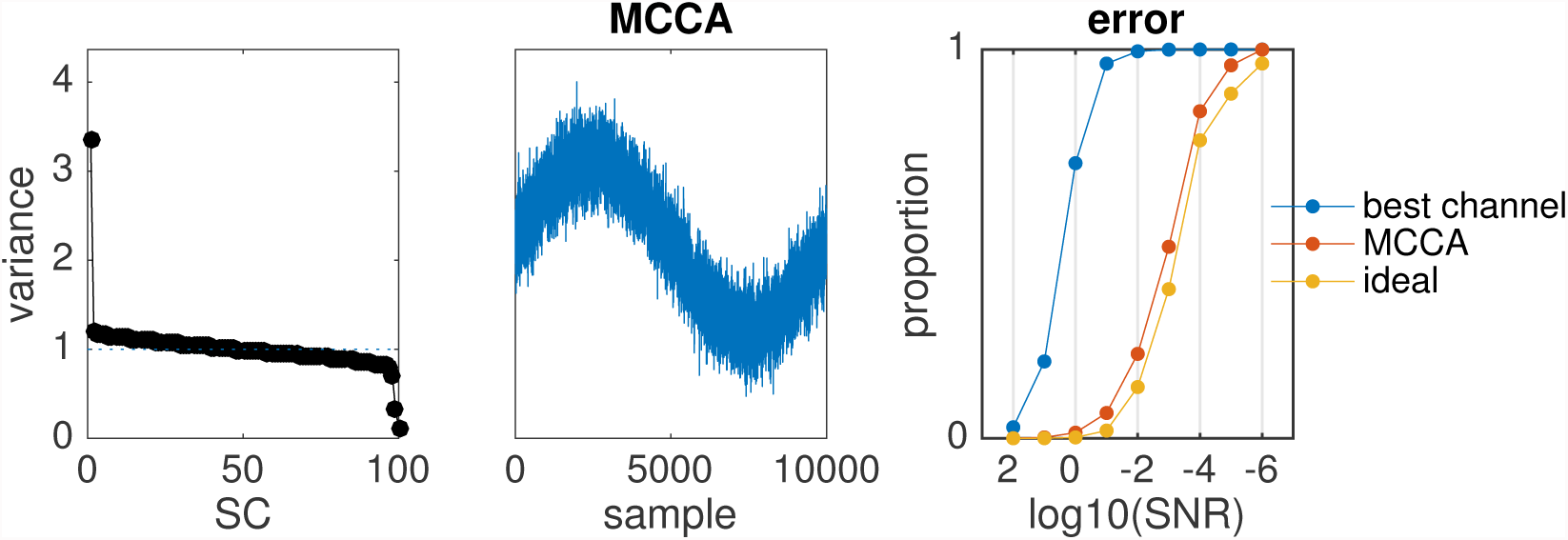
Simulation with inseparable noise. Left: variance of SCs as a function of their order at SNR=10^−2^. Center: target signal recovered from mixture at SNR=10^−2^. Right: proportion of residual noise power as a function of SNR for the raw data (blue), first SC (red) or ideal demixing matrix (yellow).

#### Example 3 - sinusoidal target in real EEG noise

EEG background noise differs from the white Gaussian noise that was used in the previous simulations in several ways: it usually has full rank (in particular because of electrode-specific noise), but the variance is unequally distributed across dimensions. It is also temporally structured, with strong temporal correlation and an overall low-pass spectrum. The first component recovered by MCCA is plotted in Fig. 6 (right) for several values of SNR. For SNRs of 0.1 or better the target is almost perfectly recovered. At SNR=0.03 the recovered waveform is somewhat noisy, and at SNR=0.01 or below the target is lost. For comparison Fig. 6 (left) shows the time course of a raw data channel (the channel that showed the largest correlation with the target). For SNR=10 the target waveform is obvious in the raw data, but for smaller values of SNR it is lost in the EEG noise. Comparing Fig. 6 left and right, there is a range of SNRs (roughly 0.03 to 1) for which MCCA provides a clear benefit. Below SNR=0.03 the algorithm switched to some other component within the data (Fig. 6 right, lowest trace) that happened to be similar across data matrices because of random correlations.

**Figure 6:**
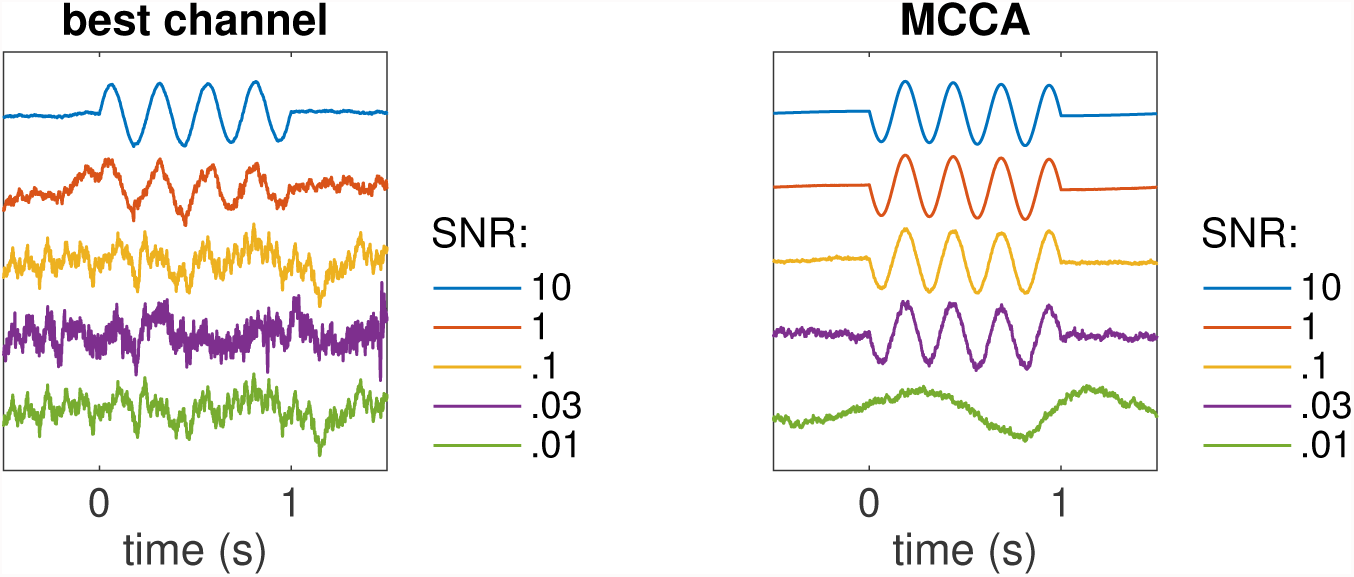
Simulation with EEG noise. Left: time course of the best raw data channel for several values of SNR. Right: time course of the first MCCA component for several values of SNR.

To summarize, MCCA is effective at extracting a weak target from within real EEG noise.

### 3.2 Real data

#### Example 4 - EEG response to tones

In this example, contrary to the previous one, the target is not known. However, since the data were collected in response to multiple repeats *and* for multiple subjects, we can apply two different methods (JD and MCCA) to isolate stimulus-evoked activity common to all subjects and compare the results. JD finds a linear transform that optimizes signal to noise ratio assuming that the signal repeats over trials. Figure 7 (top) shows the result of applying the JD analysis to the data of one subject. In the plot on the top left, the blue line shows the mean over repeats of the first component, and the gray band shows ±2 SD of a bootstrap resampling of this mean. On the top right is the topography associated with this component (computed as the map of cross-correlation coefficients between the component and each channel (Haufe et al., 2014)). MCCA can similarly be used to design a subject-specific spatial filter that improves SNR. The plots on the bottom of Figure 7 show the result of applying the subject-specific matrix derived from the MCCA analysis for the same subject. Despite the different criteria used by the two analyses (consistency over trials for JD, consistency between subjects for MCCA) the patterns are remarkably similar. To summarize, it appears that MCCA can exploit between-subject consistency to find a spatial filter that is as effective as that found by JD that exploits between-trial consistency. This is useful for data that do not involve repeated trials.

**Figure 7:**
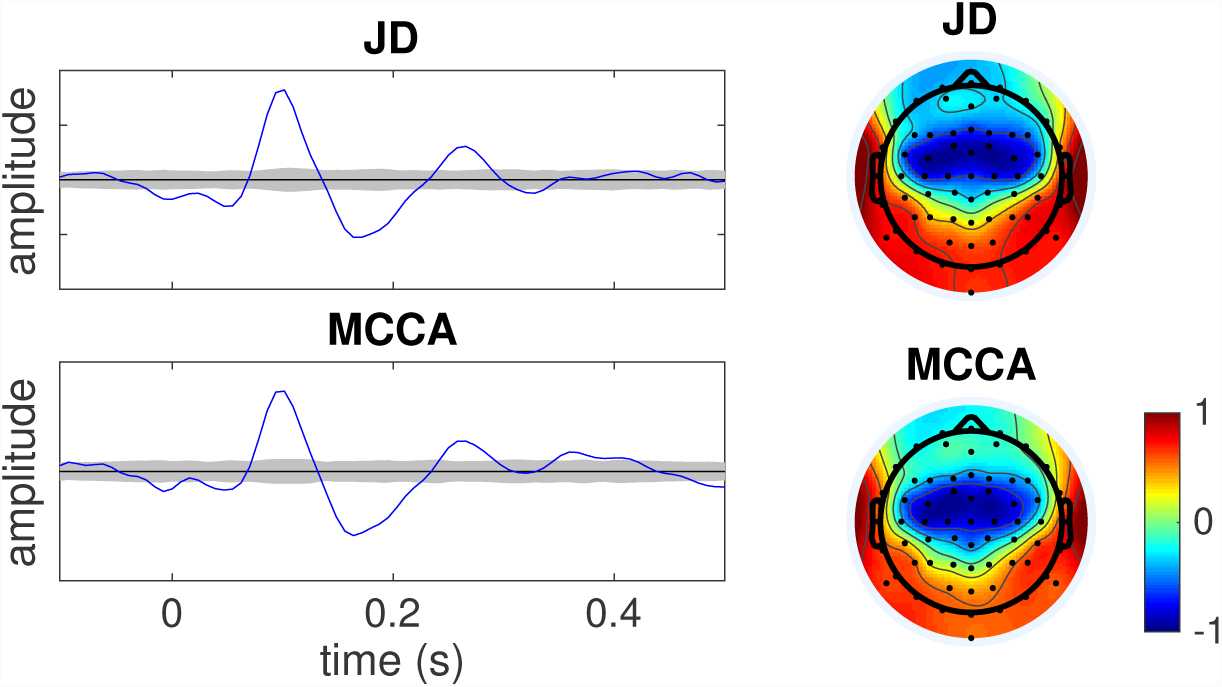
Comparison between JD solution (within-subject repeat-reliability) and MCCA solution (between-subject similarity) for one subject among ten. Data were in response to repeated tones. Left: average over trials (blue) and ±2 SD of a bootstrap resampling (gray) of the first JD component, which maximizes reliability across trials (top), or first subject-specific CC (bottom). Right: associated topographies (correlation between trial-averaged component and trial-averaged electrode waveforms).

The subject-specific MCCA analysis matrices (**V***_n_*) transform each subject’s data (**X***_n_*) into CCs (**Y***_n_*) that are well correlated across subjects so that it makes sense to average them across subjects and interpret the SCs (Y) as reflecting shared activity. Figure 8 top left shows the trial- and subject-averaged time course of the first SC, which can be interpreted as our best estimate of stimulus-evoked activity common to all subjects. It benefits from several stages of enhancement: (a) spatial filtering within each subject, (b) averaging over trials, (c) averaging across subjects. Also shown in Fig. 8 are the ten subject-specific topographies associated with this component. Despite some differences, topographies are quite similar across most subjects except S1. The bottom left plot shows the maximum over electrodes of the correlation coefficient between the first SC and each electrode (trial-averaged). Correlation coefficients are relatively high except for Subject 1 for whom the EEG response did not match the other subjects.

**Figure 8:**
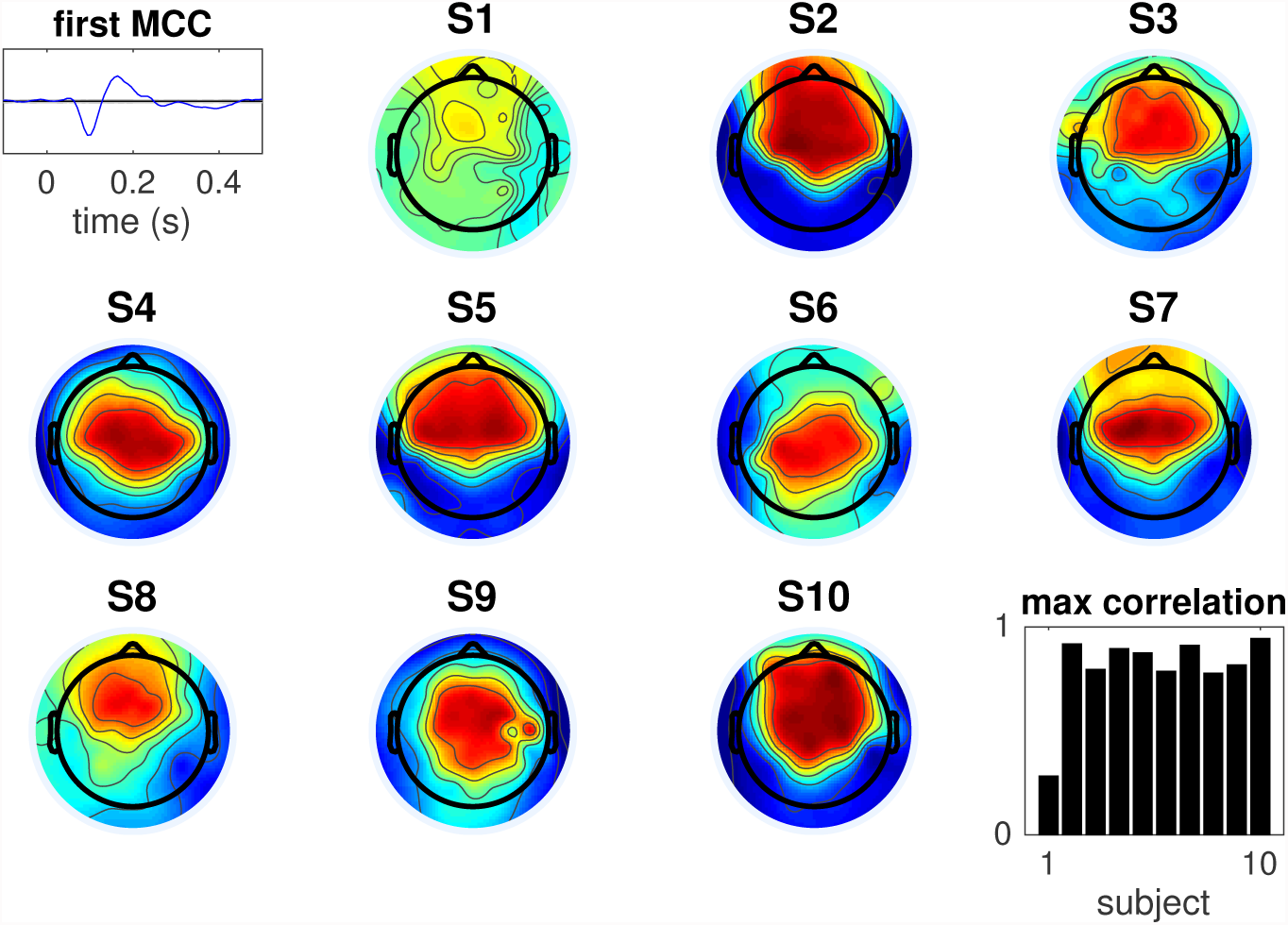
MCCA analysis of tone response, summary over 10 subjects. Top left: trial-averaged time course of the first SC. Bottom right: maximum absolute value of correlation between that component and each electrode, for each subject. Other panels: topography of correlation values (of the SC with each electrode) for each subject (the color code is the same as in Fig. 7, bottom).

#### Example 5 - EEG response to speech

For stimuli presented once only, one cannot use repetition to distinguish the brain response from the noise. Instead, systems identification techniques (Lalor et al., 2009; Holdgraf et al., 2017; Crosse et al., 2016) are used to fit an encoding model to estimate the part of brain response that is driven by the stimulus, using some representation of the stimulus (e.g. envelope or spectrogram) that can be linearly related to the brain signals. The part of the response that fits the model can be taken as the “true” response, and the rest discarded as noise. However, this partition is contingent on the validity of the stimulus representation and the quality of the model. With MCCA, a “ground truth” response can instead be estimated based on similarity of brain responses across subjects.

EEG were recorded in response to continuous speech (see Methods), and a model was fit to stimulus and response to capture their correlation (de Cheveigné et al., 2018; Dmochowski et al., 2017). The model used CCA to form pairs of maximally-correlated linear transforms of the audio stimulus features and of the EEG respectively (audio-EEG CCs). Note that this usage of CCA is unrelated to our usage of MCCA to merge data across subjects. The quality of that model was evaluated using a match vs mismatch classification task (see Methods). We compute *correlation*, *d-prime* and *percent correct* classification scores to evaluate the benefit of inserting a stage of MCCA-based denoising within the EEG preprocessing pipeline.

Figure 9 (a) shows the correlation between the first audio-EEG CC pair (thick blue line) and subsequent pairs (thin lines), with and without MCCA-based de-noising, for one subject. To the extent that correlation is limited in part by EEG noise, the higher scores on the right suggest that denoising was effective. The d-prime metric measures the degree of separation between distributions of correlation scores for matched and mismatched segments. Figure 9 (b) shows the d-prime metric for the first pair (thick blue) and subsequent pairs (thin lines), with and without MCCA-based denoising for segments of duration 64 s. The dotted line shows the d-prime metric for the multivariate distributions of audio-EEG CC pairs. The larger d-prime scores with MCCA-based denoising suggest that it can effectively contribute to improved discrimination. Figure 9 (c) shows classification scores as a function of segment duration with (red) and without (black) MCCA-based denoising. The higher scores with MCCA-based denoising show its benefit for this task. Figure 9 (d) shows that a similar benefit is found in all subjects. The thick lines are scores for a duration of 16 s, whereas the thin lines are for segments of 2 s (lowest lines) or 64 s (highest lines). To summarize, MCCA is of benefit as a denoising tool for EEG responses to speech.

**Figure 9:**
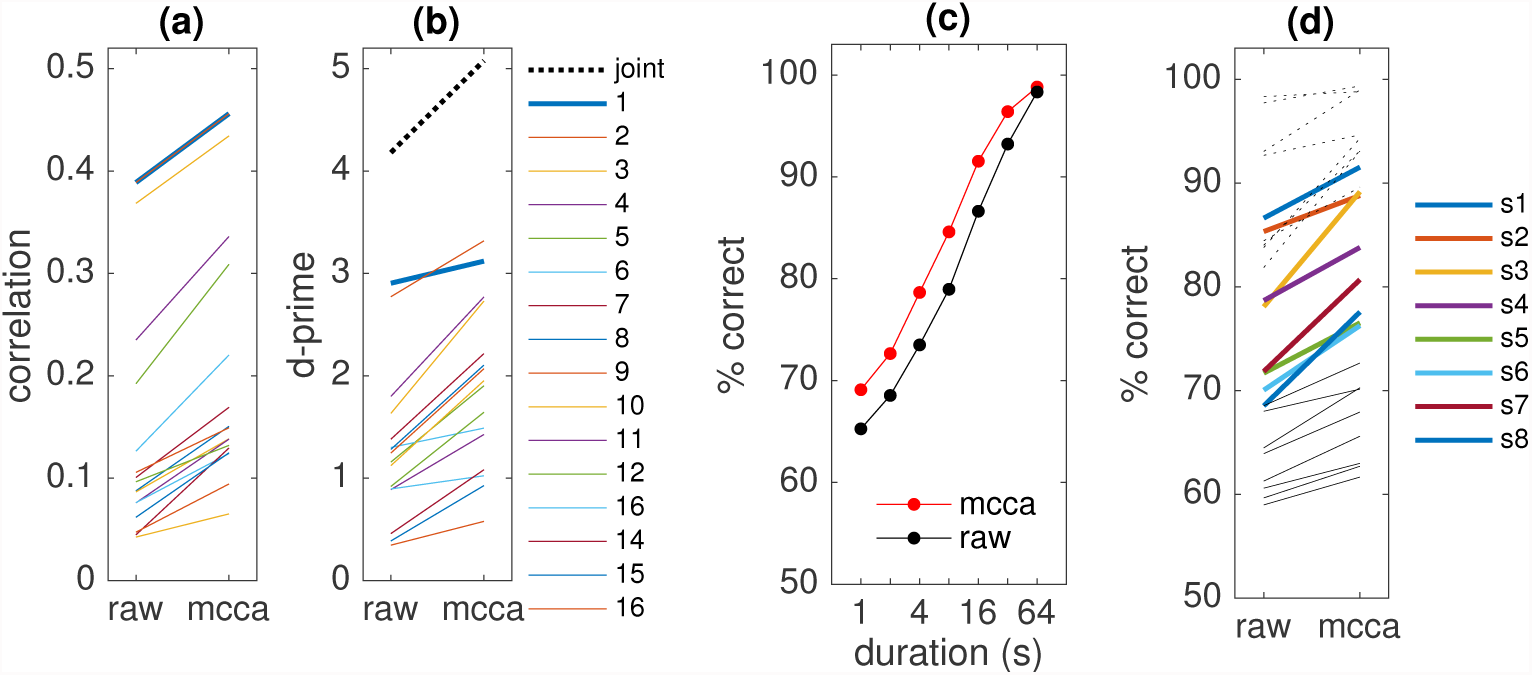
Speech-EEG decoding. (a) Correlation coefficient for the audio-EEG first CC pair (thick blue line) and subsequent pairs (thin lines) for a CCA model, with and without MCCA-based denoising. (b) d-prime metric for a classification task for the first audio-EEG CC pair (thick blue line) and subsequent pairs (thin lines), with and without MCCA-based denoising. The dotted line is for multivariate classification based on all CC pairs. (c) Percentage correct classification as a function of interval duration, with and without MCCA-based denoising. (d) Percentage correct for intervals of duration of 16s (thick lines) for 8 subjects, with and without MCCA-based denoising. Thin lines are scores for 64 s (uppermost) or 2 s (lowermost).

#### Example 6 - fMRI responses to natural sounds

Data were taken from a study that investigated fMRI responses to natural sounds (Norman-Haignere et al., 2015), in which 10 subjects listened to a set of 165 sounds belonging to 11 different classes. MCCA was applied to find patterns of selectivity to sound that were common across subjects as explained in the Methods. In brief, the 165 × 6309 matrix of voxel activations for each subject was reduced to a 165 × 12 matrix using SVD, the reduced matrices concatenated, and submitted to PCA to obtain a 165 × 120 matrix of SCs. Their variances are plotted in Fig. 10 (top left). The first 10 SCs were subjected to a JD analysis to enhance the contrast between musical sounds (classes ‘Music’ + ‘VocalMusic’) and other sounds as explained in the Methods.

**Figure 10:**
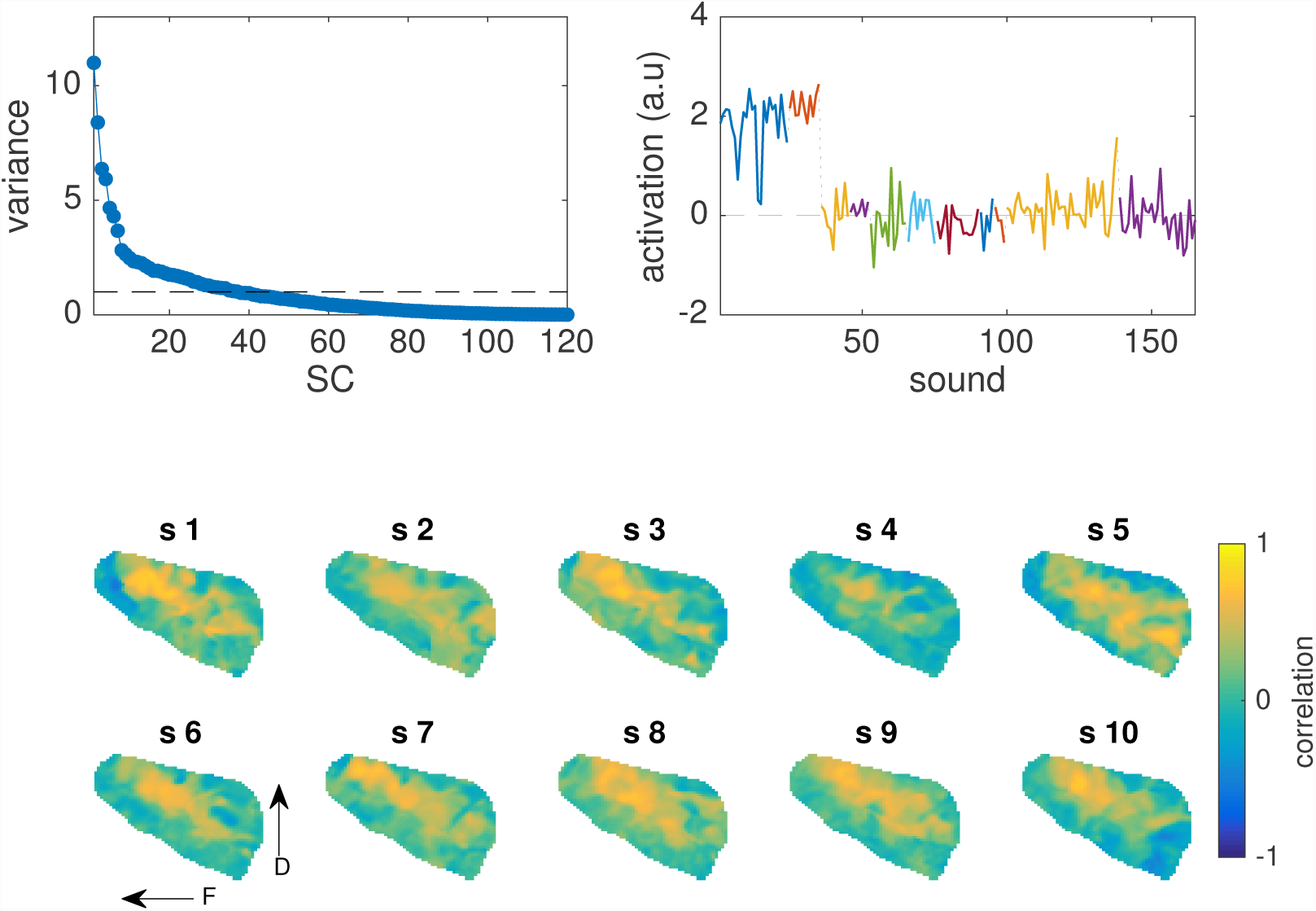
MCCA off MRI responses to natural sounds. Top left: SC variance as a function of order. Top right: activation as a function of sound of a component selective for music obtained by applying JD to the first 15 SCs (see text). Each color represents a different sound category; the first two categories are’ music’ and ‘vocal music’. Bottom: topographies of correlation between the music-selective JD component and the profile of response over sound of each voxel of the right hemisphere, for each subject.

The profile of activation over sounds of the first JD component is plotted in Fig. 10 (top right), with sounds ordered by class and coded as different colors. Activations of the first two classes (’Music’ +’VocalMusic’) are clearly distinct from that of the other classes. The corresponding topography of activation over voxels for each subject can be calculated by cross-correlating this component with the profile of activation over sounds of each voxel. Topographies for the left hemisphere for all subjects are plotted in Fig. 10 (bottom). To a first approximation, topographies are consistent in that a dorso-frontal concentration of activity is found in most subjects. To a second approximation, each topography includes additional regions, suggesting a wider network of activation that is more subject-specific. Such subject-specific details would be smoothed out by averaging over subjects. A similar JD analysis to enhance speech-specific activation revealed patterns with more ventral topographies (not shown). The outcome of this analysis is consistent with that reported by Norman-Haignere et al. (2015) using an ICA-related technique.

The benefit of MCCA here can be interpreted in terms of dimensionality reduction, based here on *consistency across subjects* rather than variance as with PCA. Dimensionality reduction allowed the final JD analysis to be performed on a matrix of size 165 × 12 × 10 rather than 165 × 6309 × 10, making it more effective by reducing overfitting. If PCA had been used instead of MCCA, the 12 selected dimensions might well have been dominated by noise. Using MCCA ensures that they are instead dominated by activity similar across subjects, which is likely to be relevant because all subjects heard the same stimuli.

This example demonstrates that MCCA can be applied also to data with more channels (pixels or voxels) than data points. MCCA offers a powerful, alternative, way of summarizing the high-dimensional data without having to explicitly model what parts of the brain response are driven by the stimulus features.

## 4 Discussion

MCCA finds a linear transform applicable to each data matrix within a data set to align them to common coordinates and reveal shared patterns. It can be used in several ways: as a *denoising* tool applicable to an individual data matrix, as a tool for *dimensionality reduction*, as a tool to *align* data matrices within a common space to allow comparisons, or as a tool to *summarize* data and reveal patterns that are general across data matrices. As formulated here, MCCA is easy to understand, straightforward to apply, and computationally cheap. Care is nonetheless required when applying it, in particular to avoid phenomena such as overfitting.

### What is new?

As reviewed in the Introduction, several versions of MCCA have been proposed in the literature and applied to the analysis of brain data. The contributions of this paper are the following. First, the formulation as a cascade of PCA, normalization, concatenation, and PCA offers an intuitive explanation that may help practitioners gain insight into this method. Past formulations may be hard to follow for the non-mathematically inclined, and their sheer number is bewildering. We used a similar 2-step formulation in a recent tutorial on joint decorrelation (de Cheveigné and Parra, 2014), and we hope that the present paper too will have tutorial value. Second, our usage of MCCA as a denoising tool, to attenuate noise within individual subjects based on across-subject consistency by projection on the overcomplete basis of its SCs, seems to be new. Third, we provide tutorial examples that may encourage researchers to put MCCA to work for a wider range of tasks, including denoising, outlier detection, summarization, and cross-subject statistics.

### How does it work?

The effect of the processing steps is schematized in Fig. 11. Multiple data matrices contain the same source component S, illustrated as a color gradient, mixed here into two 2-dimensional data matrices (Fig. 11a). Each point represents a sample in time (row of the data matrix) and the two axes represent two channels (columns of the data matrix). The color could represent a hidden sensory response that is similar across two subjects. The initial PCAs sphere each data matrix (b), so that the cloud of points is free to rotate in any direction. However, concatenating the sphered data matrices creates a cloud (in a 4-dimensional space) that is not spherical because of the shared component correlation along some direction in 4-D space (projected to 2D in panel (c)). The second PCA finds this direction of correlation between the data matrices and aligns it with the first axis (d), in the process transforming each data matrix so that it is optimally aligned with the other (e).

**Figure 11:**
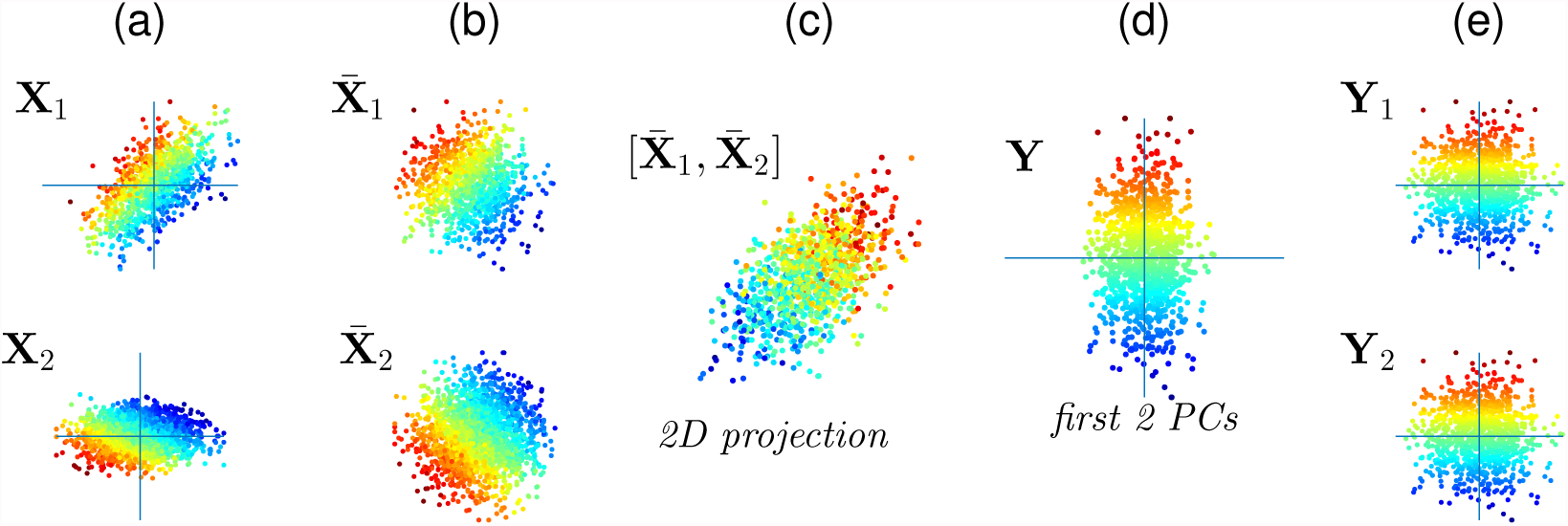
Principle of MCCA. (a) Several data matrices share a common component (coded as color) but its orientation and nature are unknown. (b) Whitening makes the data matrices free to rotate. (c) Concatenation creates a cloud in 4D space (projected here to 2D) with a direction of greater correlation/variance due to the shared component. (d) The second PCA aligns this direction with the axes. (e) In the process, the whitened data matrices are rotated such that shared dimensions are maximally aligned.

### Relation with other formulations of CCA and MCCA

As explained by Parra (2018), the aim of MCCA is to find projection vectors v*_n_* applicable to **X***_n_* that maximize the ratio of between-set to within-set covariance:

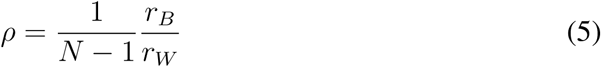

with:

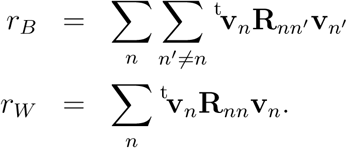

where **R**_*nn*_ = ^t^**X**_*n*_**X**_*n*_ and 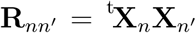, are covariance and cross-covariance matrices of the data. The divisor 1 *– N* ensures that ρ scales between 0 and 1. Setting to zero the derivative of *ρ* with respect to v*_n_*, the solution is obtained by solving the equation

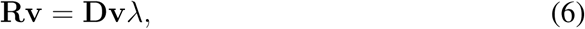

with

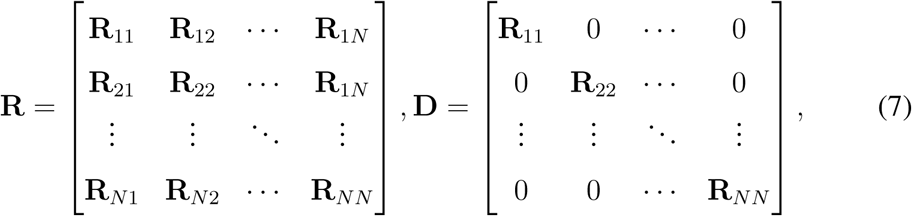

where λ = *ρ*/(*N* − 1) + 1. Now, first decompose **D** = **UΛ** ^t^**U**. Because **D** is the block-diagonal matrix of the covariances in each data set, this decomposition implies doing PCA on each data set separately, i.e whitening each data set. With this decomposition Eq. 6 can be rewritten as:

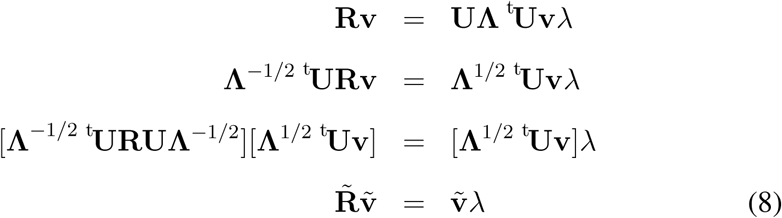

where 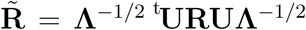 is the covariance of the whitened concatenated data. Equation 8 thus corresponds to performing PCA on the concatenated whitened data. In summary, the two-step PCA describe in the Methods (’simple MCCA formulation’) maximizes correlation between data sets. This corresponds to the standard SUMCORR formulation of MCCA described by Kettenring (1971) (see Parra, 2018). The relations between this and other MCCA formulations are described in (Asendorf, 2015).

### MCCA vs CCA

MCCA is understood as a generalization of CCA but some differences are worth noting. For CCA the focus is usually on the CCs **Y***_n_*(*n* = 1, 2), whereas for MCCA it may also be on the SCs Y. For standard CCA the projection matrices are restricted to *d* (or min*_n_ d_n_*) columns for each data set, whereas for MCCA it may be useful to consider more than *d* columns (as in Example 5). If the objective were to capture sources common to *all* data matrices, *d* components would suffice, but to capture also sources shared by *several sources but not all*, more than d columns are required. For CCA the *d* columns of **Y**_1_ are mutually uncorrelated as are those of **Y**_2_, whereas for MCCA the *D* columns of **Y***_n_* are mutually correlated in general. Columns of their sum **Y** are uncorrelated, however.

The large number (*D* > *d*) and non-orthogonality of the columns of **Y***_n_* might be disconcerting for the researcher familiar with CCA. The method may be modified such that **Y***_n_* is instead constituted of d orthogonal columns. For this, MCCA is applied as above, for each n the first column of **Y***_n_* is projected out of **X***_n_*, and MCCA applied again. This deflationary procedure terminates after d steps because the dimensionality of each data matrix is then exhausted. Smaller matrices with orthogonal columns might be convenient in certain situations, but as pointed out they might not capture all shared sources. The procedure described in the Methods is better in this respect.

### Group analysis of multi-subject data

Gathering data from multiple subjects in response to the same stimulus serves several purposes. First, to counteract variability by increasing the number of observations, analogous to recording from repeated trials. Second, to make inferences at the population level via group-level statistical analysis. Third, to allow data-dependent analysis to improve SNR based on similarity between subjects, analogous to methods that improve SNR based on similarity between trials (de Cheveigné and Parra, 2014).

The conventional strategy of calculating a “grand average”, with corresponding channels or voxels of each subject being averaged together (Choi et al., 2013; Luck, 2005), is hampered by inter-subject differences in source-to-sensor mapping. The problem is mild for sources with broad topographies (as in Fig. 8), but for sources with more local spatial characteristics a mismatch between subjects may result in destructive summation. A similar problem affects measures of inter-subject correlation (ISC) applied directly to channels or voxels (Hasson et al., 2004), or to linear combinations that assume the same mixing vectors for all subjects (Dmochowski et al., 2012; Parra et al., 2018).

One simple expedient is to select, for each subject, a group of channels based on responses to a “localizer” stimulus or task, calculate a root mean square average waveform over these channels, and then average these over subjects (e.g. Chait et al. (2010)). However, this packs the multidimensional cortical activity into a single time course from which it may be hard to infer the richer dynamics of cortical activity. Another approach is to apply inverse modeling to map the activity to a source space common across subjects (Litvak and Friston, 2008). However, this requires accurate anatomical information for each subject and is subject to the validity of the reconstruction models (Mahjoory et al., 2017), as well as between-subject variability in source positions and orientations (Lio and Boulinguez, 2016).

Data-driven methods such as MCCA are attractive in that they find a mapping between subjects based only on shared temporal aspects of the data, without requiring external information. MCCA and related methods have been widely used for fMRI data (Li et al., 2009; Correa et al., 2010b; Hwang et al., 2012; Afshin-Pour et al., 2012; Karhunen et al., 2013; Haxby et al., 2011; Afshin-Pour et al., 2014) and EEG/MEG (Lankinen et al., 2014; Sturm, 2016; Zhang et al., 2017). In contrast to MCCA, which finds variance dimensions that are similar across subjects with no attempt to ensure that they correspond to sources within the brain, ICA-based approaches attempt to to isolate sources common across subjects based on criteria of statistical independence (Calhoun and Adali, 2012; Eichele et al., 2011; Huster et al., 2015; Chen et al., 2016; Madsen et al.; Huster and Raud, 2018). Group ICA (GICA) as formulated by Eichele et al. (2011) can be seen as a concatenation of MCCA (as described here) with ICA. Isolating the MCCA step, as we do here, is useful conceptually and avoids the computational cost and assumptions associated with ICA. Hyperalignment, as used by Haxby et al. (2011), is conceptually the same as MCCA but restricting the transformations to rotations, i.e. Procrustes analysis (Xu et al., 2012). Hyperalignment has the advantage to maintain metric distance of patterns in the original and transformed space, but the disadvantage that it cannot favor channels with higher inter-subject correlation.

The focus here is on *temporal patterns* common to all subjects and thus in the MCCA procedure the data are concatenated along the spatial dimension (channels). It is also possible to extract *spatial patterns* common across subjects by concatenating data along the temporal dimension. Methods for group analysis of data from multiple subjects are reviewed by Correa et al. (2010a,b); Calhoun and Adali (2012); Sui et al. (2012); Afshin-Pour et al. (2014); Dähne et al. (2015); Chen et al. (2016); Huster and Raud (2018).

### Denoising and dimensionality reduction

As described in the Methods and illustrated in the Results, data from single subjects can be denoised by projecting on the overcomplete basis of *D* CCs, truncating, and projecting back. Data dimensions that are not shared with other subjects are *downweighted* but not removed, so in general the rank of the data remains the same. Setting the cutoff 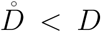 to a relatively high order suppresses only those components that are very different from those found in other subjects, most likely to be noise. In Example 5, the set of 40 PCs that represented each subject were transformed into 320 CCs, of which 110 were selected before being projected back to obtain “denoised” data, yielding the benefit shown in Fig. 9. The CCs that were rejected absorbed some of the subject-specific patterns of noise, improving the outcome.

It is often useful to reduce the dimensionality of the data for computational reasons (to reduce memory or computation time), or to avoid overfitting. The standard procedure of applying PCA and truncating the series of PCs implicitly equates variance to relevance, which may not be justified, as artifact sources may have high variance, and useful sources may be weak. MCCA is of use in this respect to replace the variance criterion by a criterion of consistency with other data. This can be done conservatively by removing a small fraction of SCs that represent the most atypical patterns within the data set.

As a tool to analyze or denoise the data of a single subject, MCCA is comparable to data-driven linear analysis techniques such as PCA, Independent Component Analysis (ICA), Joint Diagonalization, CCA and others. The fact that it uses a different criterion makes it *complementary* to those methods as a denoising or dimensionality reduction tool (e.g one can apply MCCA before or after ICA, JD, etc.).

### Caveats and cautions

A risk, common to other data-driven methods such as ICA or JD, is circularity of the analysis (Kriegeskorte et al., 2009). The method is designed to optimize correlation between data matrices, and therefore the observation that the components that it finds *are* correlated between data matrices is of little weight, unless corroborated by careful cross-validation. Related to this issue is overfitting: each SC depends on 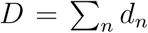 parameters, a number that can be large if there are many data matrices involved. Overfitting can be detected using resampling and cross-validation methods, and the risk of overfitting can be reduced by dimensionality reduction or other regularization techniques.

MCCA can easily latch on to artefacts and noise patterns shared across data matrices. Uninteresting linear or polynomial trends (for example EEG drift potentials) may thus appear among the first MCCA components. More generally, MCCA can be biased towards narrowband or low-frequency components common across data matrices, *even if their phase is not aligned*, particularly if the noise is spectrally-shaped or contains narrow-band components. This is illustrated in Fig. 12 that shows the result of applying MCCA to ten “data matrices”, each of 12 s duration, extracted at random from the same 40-channel EEG data that was used as background noise in Example 3. No known signal is common across these data matrices, nonetheless the lowest-order SCs have narrow spectra (Fig. 12 left) and quasi-sinusoidal waveforms (right) that might make them seem significant. It is easy to understand why MCCA might take such components to be shared: a sinusoid of arbitrary phase can be expressed as the weighted sum of a sine and a cosine, and thus narrowband activity can be approximated as resulting from two sinusoidal components in quadrature phase. As this is the case for all datasets, MCCA will select the two-component sinusoidal basis as common. Such spurious components compete with genuine shared activity, complicating the analysis.

**Figure 12:**
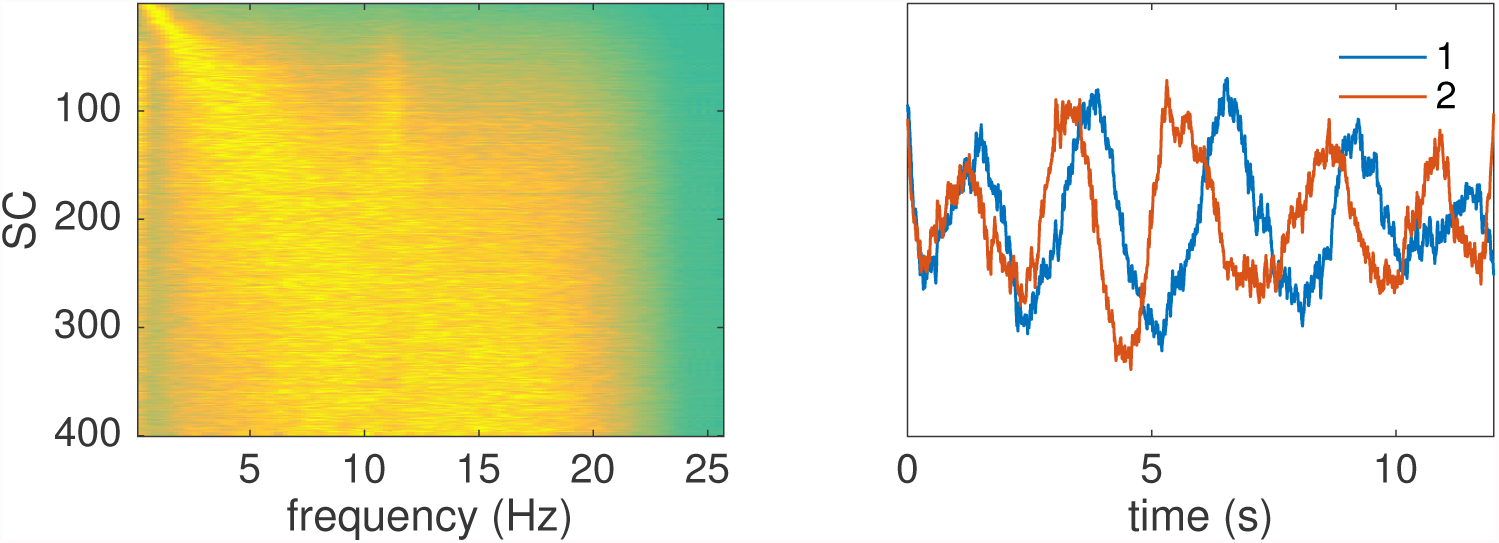
MCCA’s bias towards narrowband and low-frequency activity. Left: power spectra of SCs derived from an MCCA analysis of 10 EEG “data matrices” of duration 12 s randomly sampled from 40-channel EEG data. Power is coded as color. Right: time course of the first two SCs.

MCCA assumes that temporal patterns are common across data matrices. A difference in latency of a brain response between different subjects may reduce the ability of MCCA to extract this activity. A common outcome in that case is two components, one with a shape similar to the average pattern over subjects, and the other similar to their difference (or derivative). MCCA can readily be extended to include time-lags to account for differences in response latency between subjects, although this comes at the expense of a greater number of parameters and a greater risk of overfitting. MCCA is obviously of no benefit in the absence of synchronous patterns, for example it is not well suited for analyzing resting-state data of a group of subjects.

MCCA yields both CCs and SCs, either of which can be exploited. When reporting, it is important to specify which, to avoid confusion. As an example, the phrase ‘MCCA was applied as a preprocessing step’ is not sufficient to specify what was done.

### Applicability to real-time processing

This work was motivated in part by the need to steer an auditory assistive device using brain signals. An obstacle to reliable decoding is the high-level of noise and artifacts in the EEG signals, and analysis and denoising methods are essential for the success of this application. To be useful, a method must be applicable to *real-time* processing, whereas MCCA as described here works in batch mode. It may nonetheless be of use in the following fashion. EEG data is recorded from a pool of subjects to a calibration sample of speech, and MCCA is used to derive a “canonical” EEG response to that sample. To adapt the system to a new user, EEG data are recorded in response to the calibration sample, and a spatial filter is designed (for example using CCA) to maximize similarity between the subject’s and the canonical response. This spatial filter is then used in the real-time processing pipeline. This suggests that MCCA can also be put to use in a practical application such as cognitive control of a hearing aid.

## 5 Conclusion

Multiway CCA is a powerful tool for analysis of multi-subject multivariate datasets. It can be used both to design spatial filters to denoise data of each individual subject, and to summarize data across subjects. Many related methods have been proposed in the literature, but the processing principles behind them, and the range of tasks that they can be used for, are not widely appreciated. The use of MCCA (or similar techniques) should be more prevalent given the ubiquitous need for merging data across subjects. In this paper we presented a formulation of MCCA that is relatively easy to understand, illustrated in detail how it works, and showed how it can be put to use for a wide range of common tasks in multi-subject multivariate data analysis.

## Acknowledgements

This work was supported by the EU H2020-ICT grant 644732 (COCOHA), and grants ANR-10-LABX-0087 IEC and ANR-10-IDEX-0001-02 PSL*. Lucas C. Parra received support from the National Science Foundation under grant DRL-1660548. Some of these ideas were tried out at the 2017 Telluride Neuromorphic Engineering Workshop. Malcolm Slaney and Sam Norman-Haignière offered useful comments on earlier versions of the manuscript.

